# Progress in the molecular Phylogeny of *Cotesia acuminata* and *C. melitaearum* cryptic species complexes

**DOI:** 10.1101/2025.08.19.670998

**Authors:** Julian Schach, Federica Valerio, Anne Duplouy

## Abstract

Cryptic species present major challenges for biodiversity and evolutionary research due to the morphological similarity and frequent lack of genetic and ecological data. In parasitoids wasps, this is especially true for lineages with limited sampling and unclear species boundaries. Here we reconstruct the phylogeny of *Cotesia* wasps parasitizing Melitaeini butterflies, including two cryptic species complexes (*Cotesia acuminata agg.* and *C. melitaearum agg.*). Using a ten-gene dataset from samples collected in Europe, Asia, and North America, we inferred relationships among 22 *Cotesia* species using maximum likelihood. We also included non-Melitaeini associated *Cotesia* to assess whether the Melitaeini parasitoids form a monophyletic group. Our analyses yielded a highly supported phylogeny, revealing four major clades, three of which included the Melitaeini associated species. This confirms that the *Cotesia* attacking Melitaeini butterflies are polyphyletic, resulting from independent host shifts across the genus. Each clade is further subdivided into subclades corresponding to the different (cryptic) species, clarifying previously unresolved relationships. These results provide a robust framework for future studies on the evolution, ecology, and host use dynamics of *Cotesia* wasps and highlight the utility of multi-locus data for resolving phylogenies in morphologically cryptic taxa.

## Introduction

Species have historically been described using morphological data, and species relationships based on degrees of similarities of morphological characteristics. More recently, the advent of molecular techniques has refined many taxonomic/phylogenetic classifications, by allowing us to study species relationships and biodiversity with much higher resolution (Kjer et al. 2016). Cryptic species, are two or more molecularly distinct species, often accompanied by rapid speciation, which are or have been erroneously classified as a single species because they are morphologically indistinguishable (Bickford et al. 2007). Such cryptic species are known to span the tree of life (Hebert et al. 2004, Grundt et al. 2006, Kon et al. 2007, Shin and Allmon 2023). Their hidden genetic diversity has however challenged our ability to estimate the true biodiversity richness on Earth, as many remain to be discovered and characterized (Li and Wiens 2023). Furthermore, with the current biodiversity crisis, we are likely losing many such species at alarming rates even before they have been taxonomically characterized, or before their ecosystem roles were understood (Koh et al. 2004).

Parasitoid wasps make up one of the most diverse animal groups in the world (Forbes et al. 2018). They lay their eggs within or on their hosts’ body, and the parasitoid larvae develop feeding on the host, until death of the host, and emergence of the adult wasp (Whitfield et al. 2018). These parasitoids can be very efficient at killing their hosts and can act as key evolutionary pressures on their host species population growth (Cornell and Hawkins 1994) and host population dynamics (Lei and Hanski 1997). For example, Lei and Hanski (1997) showed that the specialist species *Cotesia melitaearum* parasitizes only 10% of *Melitaea cinxia* caterpillars and can still cause localized extinctions within the butterfly host metapopulation. Because of their narrow host ranges and highly efficient killing of their hosts, several species of parasitoids have become popular biocontrol agents in sustainable agriculture, contributing significantly to the $417 billion USD global value of insect biocontrol (Brock et al. 2021). For example, *C. ruficrus* parasitizes the fall armyworm (*Spodoptera frugiperda*), which is contributing to food insecurity in Africa, Asia, and the Pacific (Li et al. 2024)

Most of the worlds parasitoid diversity occurs within the superfamily Ichneumonoidea, which includes the two families Ichneumonidae and Braconidae, with approximately 60,000 described species, and many more estimated (Quicke 2014). Within the Braconidae, the subfamily Microgastrinae is the most diverse with the genus *Cotesia* Cameron (Hymenoptera: Braconidae: Microgastrinae) alone including over 300 named species (Fernandez-Triana et al. 2020), and an estimated 1500-2000 species worldwide (Mason 1981). The *Cotesia* wasps usually exhibit narrow host ranges, and closely related species often differ in their host species preferences, attacking one to few closely related lepidopteran host species (Kankare et al. 2005b). For example, although previous studies suggested that seven species of *Cotesia* were generalist parasites of the Melitaeini butterfly species found across Europe and Asia (Wahlberg et al. 2001, Kankare and Shaw 2004), more recent molecular analyses accompanied by rearing experiments provided evidence that the originally considered generalist parasitoid species, *C. acuminata* and *C. melitaearum*, were in fact numerous cryptic species, each with narrow Melitaeini butterfly host specificity (Kankare et al. 2005a, 2005b).

In the complex species community of Melitaeini butterflies occurring in Catalonia (Catalan Butterfly Monitoring Scheme, CMBS) in northeastern Spain, Kankare et al. (2005b) indeed identified that the 10 Melitaeini butterfly species found, are hosts to at least 10 *Cotesia* (cryptic) species. Unfortunately, the original barriers to geneflow between the current cryptic lineages of the wasps remain unclear, as studies exploring whether isolation by distance or endosymbiotic bacteria failed to explain the sympatric speciation of the parasitoid species (Kankare et al. 2005b, Valerio et al. 2023, Schach 2024) Currently, the phylogeny of the Catalonian *C. acuminata* complex and *C. melitaearum* complex are based on few microsatellites (Kankare et al. 2005b), while a wider Eurasian phylogeny of these groups combined two mitochondrial DNA (mtDNA) loci and a nuclear DNA (nDNA) locus (Kankare and Shaw 2004). As a consequence of the restricted available genomic material included in these phylogenies, the relationships between Cotesia species still contain several unresolved nodes (i.e., nodes with low bootstrap support estimates; Kankare and Shaw 2004, Kankare et al. 2005b). This challenges our ability to further explain the true diversity of and speciation scenarios between *Cotesia* wasps attacking Melitaeini butterflies.

Here, we aimed to refine the phylogeny of *Cotesia* species associated with 16 species of Melitaeini butterflies, by using a larger number of nuclear markers and a wider sample diversity than previous studies. We analysed sequences of two mtDNA and eight nDNA markers from both freshly collected specimens from Finland, Italy, Spain, and Switzerland, as well as DNA extracts from samples collected between 1979-2002 from China, Finland, France, Hungary, Russia, Spain, Sweden, and USA. With this, we expected to resolve the previously unclear nodes in the *Cotesia* phylogeny, particularly the basal nodes, and to provide a strong phylogenetic reconstruction within the cryptic species complexes.

## Materials and Methods

### Samples

This study includes *Cotesia* parasitoid wasp specimens that had emerged from the caterpillars of 16 species of Melitaeini butterflies. Including *Euphydryas aurinia, Euphydryas aurinia davidii, Euphydryas desfontaini, Euphydryas editha, Euphydryas maturna, Euphydryas phaeton, Melitaea athalia, Melitaea cinxia, Melitaea cynthia, Melitaea deione, Melitaea didyma, Melitaea latonigena, Melitaea parthenoides, Melitaea phoebe, Melitaea scotosia,* and *Melitaea trivia* (Table 1).

**Table 1.** *Cotesia* identity, host species, number of individuals, caterpillar host plant, country, locality, collector ID, and finally collection year. “-” indicates the information was unavailable.

| **Cotesia species** | **Butterfly host species** | **# of Ind.** | **Identified host plant** | **Country** | **Locality** | **Collector** | **Collection year** |
| --- | --- | --- | --- | --- | --- | --- | --- |
| *C. acuminata* Sp. A | *Melitaea latonigena* | 3 | - | Russia | Siberia | N.W. I.H. | 1998 |
| *C. acuminata* Sp. A | *Melitaea didyma* | 5 | *Plantago lanceolata* | Spain | Coll d'Estenalles | F.V. | 2023 |
| *C. acuminata* Sp. A | *Melitaea didyma* | 5 | - | Switzerland | Conthey | Y.C. | 2023 |
| *C. acuminata* Sp. B | *Melitaea scotosia* | 3 | - | China | Peking | I.H., L.G. | 1999 |
| *C. acuminata* Sp. B | *Melitaea phoebe* | 5 | - | Italy | Aosta | Y.C. | 2023 |
| *C. acuminata* Sp. K | *Euphydryas maturna* | 1 | - | France, | Cote d'Or, Malloy, Lindesberg | P.J.C.R. | 1999 |
| *C. acuminata* Sp. K | *Euphydryas maturna* | 3 | - | Sweden | Lindesberg | C.U.E. | 1999 |
| *C. acuminata* Sp. L | *Melitaea athalia* | 1 | - | France | Var, St. Paul-en-Foret | P.W.C. | 1982 |
| *C. bignelli* | *Euphydryas aurinia* | 5 | - | Switzerland | La Vraconnaz | Y.C. | 2023 |
| *C. cynthiae* | *Melitaea Cynthia* | 4 | - | France | Alps, Laus de Cervieres | M.S. | 2001 |
| *C. euphydryidis* | *Euphydryas phaeton* | 7 | - | USA | Warren Co., Front Royal | N.S. | 1979 |
| *C. koebelei* | *Euphydryas editha* | 7 | - | USA | Yosemite National Park | M.S., B.W. | 2001 |
| *C. melitaeorum* Sp. D | *Euphydryas desfontainii* | 5 | *Cephalaria leucantha* | Spain | Fígols-El Forn | CS | 2023 |
| *C. melitaearum* Sp. *D* | *Euphydryas aurinia* | 5 | *Lonicera implexa* | Spain | Cornudella de Montsant | C.S. | 2023 |
| *C. melitaearum* Sp. *D* | *Euphydryas aurinia* | 5 | *Lonicera implexa* | Spain | La Mussara | C.S. | 2023 |
| *C. melitaearum* Sp. *E* | *Melitaea deione* | 5 | *Antirrhinum majus* | Spain | Les Illes | CS | 2023 |
| *C. melitaearum* Sp. *E* | *Melitaea deione* | 5 | *Antirrhinum majus* | Spain | Poboleda | CS | 2023 |
| *C. melitaearum* Sp. *F* | *Melitaea didyma* | 3 | - | Hungary | Örseg | M.R.S. | 2001 |
| *C. melitaearum* Sp. *F* | *Melitaea didyma* | 2 | - | Spain | Cantallops | C.S. | 2002 |
| *C. melitaearum* Sp. *G* | *Melitaea trivia* | 10 | *Verbascum pulverulentum* | Spain | Pyrenees | C.S. | 2023 |
| *C. melitaearum* Sp. *H* | *Melitaea cinxia* | 8 | *Plantago lanceolata* | Finland | Åland | S.I. | 2022 |
| *C. melitaearum* Sp. *I* | *Melitaea athalia* | 5 | - | Finland | Åland | L.G., S.vN. | 1997 |
| *C. melitaearum* Sp. *M* | *Euphydryas aurinia davidi* | 3 | - | Russia | Siberia | N.W., I.H. | 1998 |
| *C. melitaearum* Sp. *N* | *Melitaea parthenoides* | 5 | *Plantago media* | Spain | El Brull | C.S. | 2023 |

Collectors: Y.C., Yannick Chitarro; P.W.C, Peter W. Cribb; C.U.E., Clae U. Eliasson; L.G., Lei Guanchung; I.H., Ilkka Hanski; S.I., Suvi Ikonen; P.J.C.R., Peter J. C. Russel; M.R.S., Mark R. Shaw; M.S., Michael Singer; N.S., Nancy Stamp; C.S., Constanti Stefansescu; S. vN., Saskya van Nouhys; N.W. Niklas Wahlberg; B.W., Brian Wee; F.V. Federica Valerio

Among the *Cotesia* wasps, two cryptic species complexes are known to consist of multiple species: *C. acuminata* and *C. melitaearum*. Following the convention introduced by Kankare et al. (2005a), we refer to these as *C. acuminata agg.* and *C. melitaearum agg.,* with individual lineages denoted by species-level designations (e.g. *Sp. A., Sp. B.*)

The Melitaeini caterpillars were collected over two time periods: 1979-2002 and 2022-2023. The DNA from early samples collected for the study of *Cotesia* wasps’ phylogeny and butterfly-host specialization in earlier studies (Kankare and Shaw 2004, Kankare et al. 2005a, 2005b) was extracted in the early 2000’s and preserved in the -20 °C freezers at the University of Helsinki. These DNA extracts represent 10 species from eight different countries, including China (N=3 from 1 species), Finland (N=5 from 1 species), France (N=7 from 3 species), Hungary (N=3 from 1 species), Russia (N=6 from 2 species), Spain (N=2 from 1 species), Sweden (N=3 from 1 species), and USA (N=14 from 2 species) (Table 1). Additionally, caterpillars from one locality in Finland (Åland) were also collected in the fall of 2022. Finally, parasitized Melitaeini caterpillars were collected in early April 2023 from five localities in Catalonia, Spain (Coll d’Estenalles, El Brull, Fígols-El Forn, Pyrenees), two localities in Switzerland (Vaud and Valais), and one locality in Italy (Aosta). After collection, the caterpillars were reared in the laboratory on excess of their respective host plant, until the adult wasps emerged. Freshly emerged adult wasps were killed by placing them in a freezer for 24 hours. The samples were stored in 90% ethanol at -20℃ until further manipulated.

To test whether the *Cotesia* species parasitizing Melitaeini butterflies form a monophyletic group, we also included gene sequences retrieved from *C. chilonis, C. congregata, C. glomerata, C. ruficrus, C. typhae,* and *C. vestalis,* which full genomes are publicly available from the NCBI database. These *Cotesia* species naturally parasitize species of pest moths (e.g. *Spodoptera*, or *Manduca*) or *Peiris* butterflies (Blažytė-Čereškienė et al. 2022, Miles et al. 2023). We also included respective gene sequences from two Microgastrine species (*Microplitis demolitor,* and *Microplitis mediator*), which full genomes are publicly available from the NCBI database, to use as outgroup in the phylogenies described below. Our sequences were blasted against the respective genomes to extract the sequences of the genes used in the phylogeny. All NCBI references can be found in Table 2.

**Table 2.** Additional species added to the ingroup (seven *Cotesia*) and outgroup (two *Microplitis*). Includes the host species, GenBank accession, and reference. “-” indicates the information was unavailable.

| **Cotesia Species** | **Host** | **GenBank accession #** | **Reference** |
| --- | --- | --- | --- |
| *Cotesia chilonis* | *Chilo suppressalis* | GCA_018835575.1 | (Ye et al. 2020) |
| *Cotesia congregata* | *Manduca sexta* | GCA_905319865.3 | (Gauthier et al. 2021) |
| *Cotesia glomerata* | - | GCA_020080835.1 | (Pinto et al. 2021) |
| *Cotesia ruficrus* | *Mythimna separata* | GCA_048537615.1 | - |
| *Cotesia typhae* | *Sesamia nonagrioides* | GCA_013202065.2 | (Muller et al. 2021) |
| *Cotesia vestalis* | *Plutella xylostella* | GCA_000956155.1 | - |
| *Microplitis demolitor* | *Pseudoplusia includens* | GCA_026212275.2 | (Burke and Strand 2012) |
| *Microplitis mediator* | - | GCA_029852145.1 | - |

Depending on availability, we selected one to seven parasitoid specimens per caterpillar species, and per locality sampled for molecular work. The cryptic nature of the parasitoids, and the fact that some caterpillar species can host multiple species of parasitoids makes their identification difficult using only morphology. The species identity of the parasitoids collected in the early 2000’s was previously confirmed through the *COI* and host associations (Kankare and Shaw 2004). For the samples collected in 2022 and 2024, their species identities were assigned after blasting their respective *COI* sequences against the NCBI database, and by considering their host associations, following the protocol used by Kankare and Shaw (2004)

### DNA Extraction

For the samples collected in the early 2000’s the DNA was extracted using the NucleoSpin Tissue Kit (Macherey-Nagel) following the manufacturers protocol and eluted in 50 μL of MilliQ water (Kankare and Shaw 2004). The samples have since been stored in the freezer (-20℃) at the University of Helsinki, Finland. For the samples collected in 2022 and 2023, the DNA was extracted from each whole insects, except in two cases where we pooled adult wasps that emerged from the same host caterpillar. The Macherey-Nagel NucleoSpin Tissue Kit (Düren, Germany) was used following the manufacturer’s protocol. The DNA was eluted twice in 40 μL of elution buffer to increase final yield and concentration. We do not believe the slight differences between extraction protocols affected our final results.

#### Gene selection

Genetic markers and primer pairs designed and tested by previous studies, were first selected by reviewing the available literature on *Cotesia* and Braconidae phylogenies (See Table 2. for references). All primer pairs were tested for specificity to *Cotesia* using Primer-Blast (Ye et al. 2012). This resulted in 9 usable primer pairs from the literature. Additionally, 16 BUSCO genes were randomly selected from the *C. congregata* genome (GCA_905319865.3) using BUSCO v 5.7.1 (Manni et al. 2021). Using Primer-Blast (Ye et al. 2012), we designed an additional set of 16 primer pairs to amplify these 16 BUSCO genes. The best pairs were selected based on the lowest self-complimentary score given by Primer-Blast. In total two mtDNA markers and 23 nDNA markers were tested, but only 2+8 were successfully used in this study, 13 other nDNA primer pairs designed failed to amplify in gradient PCR (Table S5). All primer pairs, from the literature and designed for this study, were designed to amplify genetic regions with a length ranging from 150 to 850 bp. All primer pairs used in the present study are listed in Table 3.

**Table 3.**
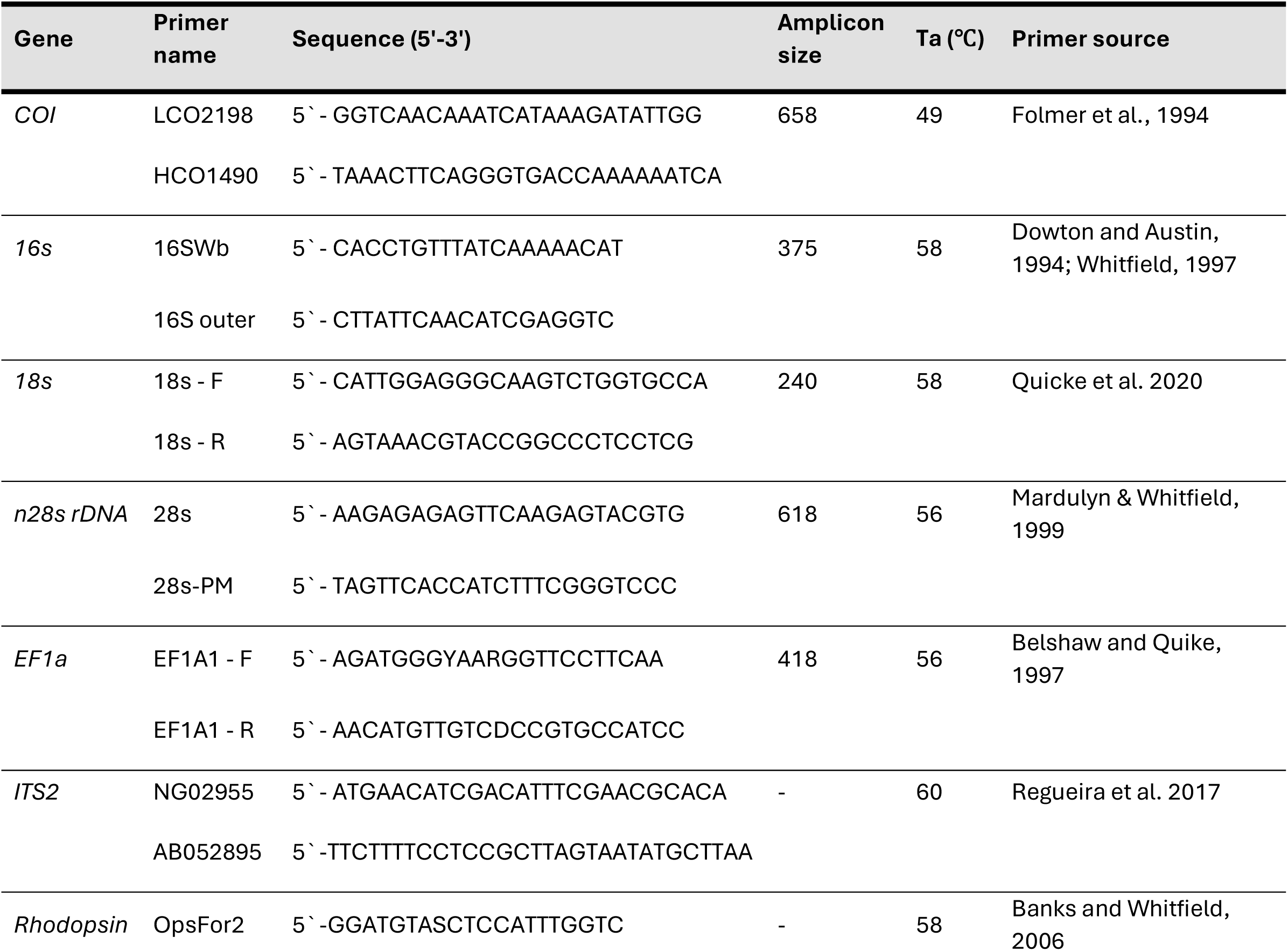
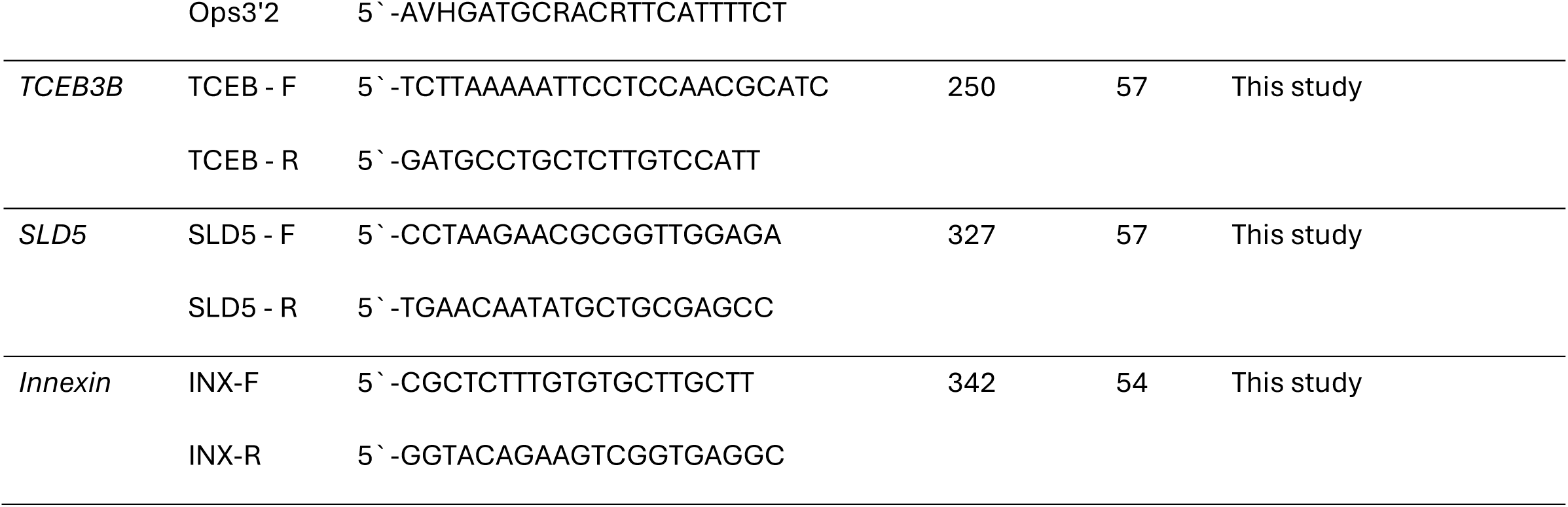
Characteristics of the primer pairs used in this study. Includes the marker targeted (gene), primer name, primer sequence, amplicon size, annealing temperature, and source. “-” indicates the information was unavailable.

#### PCR

Each 10 μL PCR reaction included of 1 μL of Buffer solution, 0.8 μL of DNTPs (25nM) mix, 0.5 μL of forward primers (10nM), 0.5 μL of reverse primers (10nM), 0.5 μL Mg2 reagent, 0.05 μL KAPA Taq polymerase, 6.7 μL of H20 (sterile), and 1 μL of DNA extract. The PCR reactions were initiated by incubation at 95°C for 3 min, followed by 30 cycles of 95°C for 40 s, the optimal annealing temperature of the respective primer pair (Table 2) for 40 s, and 72°C for 40 s. The PCR reaction ended with a final elongation step at 72°C for 5 min, and a final hold at 18°C.

The PCR products were run through a 1% agarose gel electrophoresis, with SYBR-safe stain (Thermo Scientific, USA), and visualized under UV light. For each gel, 5 μL of 100 bp DNA ladder (#SM0321, Thermo Scientific, USA) was loaded into the first well, and each subsequent well was loaded with a mixture of 4 μL of both loading dye (bromophenol blue) and PCR product. The gels were run for 30 minutes at 100 volts. The PCR amplification was repeated once for any failed sample, to identify potential false negative results.

#### Preparation for Sanger Sequencing

The successfully amplified PCR products were individually cleaned using Exonuclease I (ExoI) to remove any single-stranded DNA (Thermo Scientific, USA). For each reaction 0.5 μL of ExoI was added to 5 μL of PCR product, then the samples were incubated at 37°C for 15 minutes, followed by 15 minutes at 85°C to terminate the reaction.

Cleaned PCR products were sent to Macrogen Europe for Sanger Sequencing using the company Multiple Primer/Plate option.

#### Sequence Cleaning and Alignments

The chromatograms of each sequence were manually quality checked by using Geneious Prime 2025.01 (https://www.geneious.com). Low-quality sequences (error probability limit 0.05) and highly incomplete sequences (less than half of target length) were removed from further analysis.

We included the respective gene sequences from two Microgastrine species (*Microplitis demolitor,* and *Microplitis mediator*), to use as outgroup in the phylogenies described below. Additionally, six species of *Cotesia* that do not parasitize Melitaeini were added to the ingroup. For each maker, all sequences were aligned with MUSCLE (Edgar 2004), and all alignments were manually curated with AliView (Larsson 2014). All sequences will be uploaded to the NCBI database and accession numbers will be provided upon submission.

#### *Cotesia* Phylogeny

For the phylogenetic analysis we used IQ-TREE V2.4.0 (Minh et al. 2020). Phylogenetic analyses were conducted for: (I)all ten genes concatenated (mtDNA and nDNA), (II) eight concatenated nuclear genes, (III) two concatenated mitochondrial genes, and (IV) all genes independently. Specimens that amplified for less than half of the markers were excluded from concatenated analysis but kept for the individual gene tree analyses. We used ModelFinder (Kalyaanamoorthy et al. 2017), and the Bayesian Information Criterion (BIC) to select for the best models. The partition finding algorithm (Chernomor et al. 2016) was run for the concatenated data (ten-genes, nDNA, mtDNA) partitioned by genes using the MFP+MERGE option. Support for nodes was evaluated with 1000 ultrafast bootstrap (UFBoot) approximations (Hoang et al. 2018). Phylogenetic trees were visualized using ITOL (Letunic and Bork 2024).

## Results

### Parasitoid Identification

Many of the 2000’s samples failed the *COI* amplification and sequencing. Fortunately, those exact same samples were previously sequenced as part of earlier studies (Kankare and Shaw 2004, Kankare et al. 2005b). These N=42 specimens include10 species of *Cotesia*: four cryptic species of *C. acuminata agg.* (two of which are the same species as the 2022-2023 collections but from different hosts), *C. cynthiae, C. euphrydryidis, C. koebelei,* and three cryptic species of *C. melitaearum.* The sequencing results of the 2022 and 2023 collections confirmed that these N=68 specimens include eight species of *Cotesia*: *C. bignelli,* two species of *C. acuminata agg.,* and five species of *C. melitaearum agg*.

### Sequencing results

Of 110 samples, 41% were successfully sequenced for all 10 genes, 66% for at least 9 genes, 75% for at least 8 genes, 85% for at least 7 genes, 88% for at least 6 genes, and 90% for at least 5 genes.

### *Cotesia* Phylogenies

#### (I) Ten-gene dataset

The ten-gene dataset included 110 wasp specimens. The ten-gene alignment length was 3486 bp long, of which 1498 sites (43%) represented unique site-patterns. Out of these 747 sites (21% of all sites) were parsimony-informative. The model partitioning algorithm revealed five partitions in the ten-gene dataset. (Table S1.)

The ML analysis revealed four major clades with high support (bootstrap = 100, 77, 93) and suggested that the species attacking Melitaeini butterflies are polyphyletic (Figure 1).

- **Clade A** contains the *C. acuminata agg., C. bignelli,* and *C. euphydryidis* parasitoids from Melitaeini butterflies*. Cotesia euphydryidis* is placed at the base of the clade. The next division is between *C. bignelli* and *C. acuminata agg.* The clade reveals that *C. acuminata agg.* is monophyletic, where *Sp. B* and *K* cluster together and are sister taxa to *Sp. A.* The divergence between the *Sp. A*, and *B/K* is poorly supported (bootstrap = 49), but the divergence between *Sp. B* and *K* is well supported (Figure 1).
- **Clade B** contains *C. koebelei, C. congregata* and *C. glomerata* parasitoid wasps. *Cotesia congregata* and *C. glomerata* do not use Melitaeini butterflies as hosts but *C. koebelei* wasps, parasitize the Melitaeini species *E. editha.* The nodes in this clade are highly supported (bootstrap = 98 and above) (Figure 1).
- **Clade C** contains *C. chilonis, C. ruficrus, C. typhae,* and *C. vestalis* parasitoids, which all parasitize non-Melitaeini hosts. The divergence between these species is highly supported (bootstrap = 93 and above) (Figure 1).
- **Clade D** contains *C. cynthiae* and *C. melitaearum agg,* which parasitize Melitaeini butterflies. Ten subclades are revealed, represented by the individual cryptic species and *C. cynthiae* with moderate to high support (bootstrap 77 to 100). Here, *C. melitaearum agg.* forms a paraphyletic group. Most of the individual species within this clade group together, forming monophyletic groups, except *C. melitaearum Sp. F*. For this particular species, our phylogeny places the specimens from Spain in a sister group to *C. melitaearum Sp. G* also from Spain and the specimens from Hungary are in a sister-group to the *C. melitaearum Sp. H* from Finland. (Figure 1).

**Figure 1.**
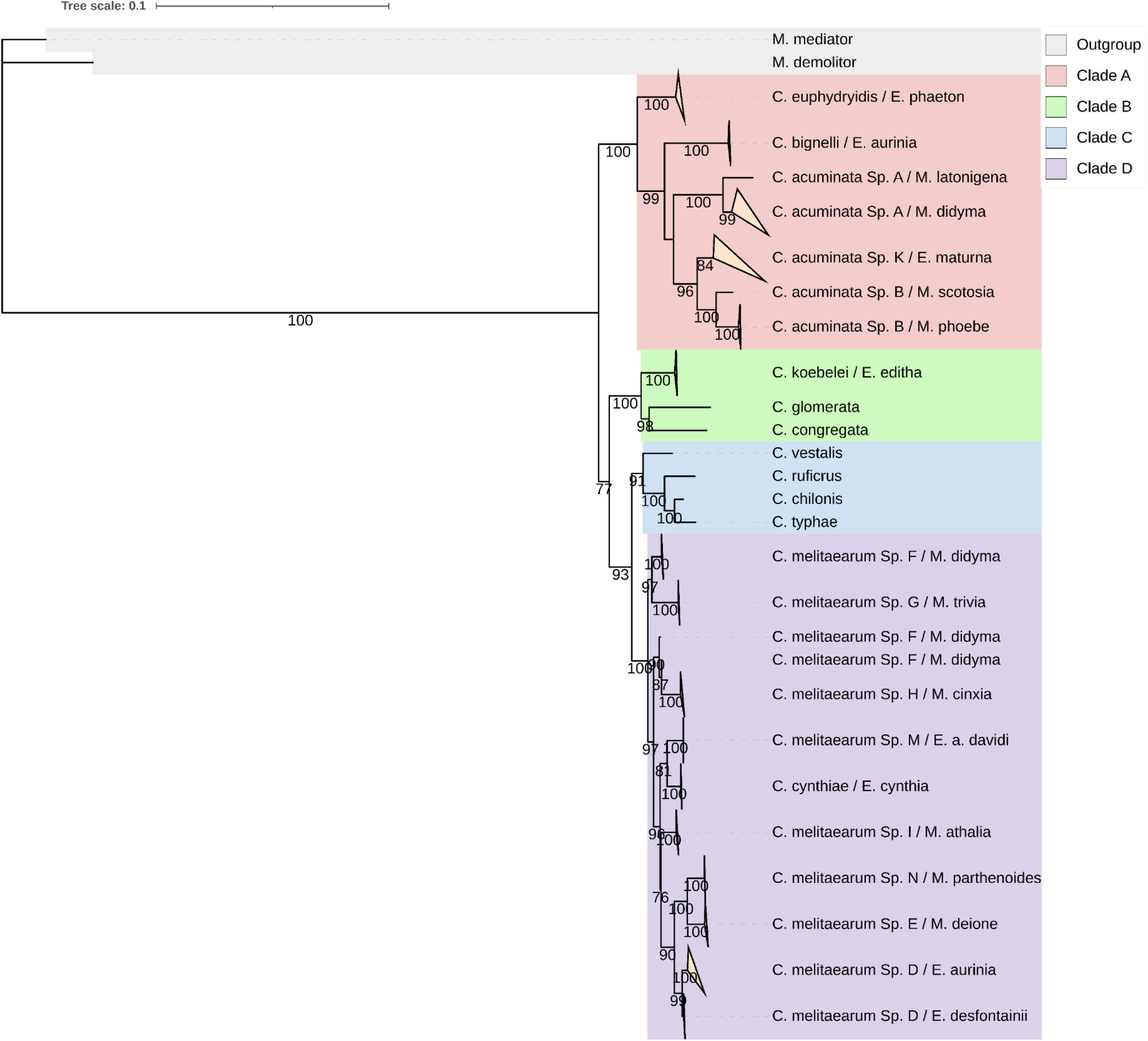
Maximum likelihood phylogeny of *Cotesia* species parasitizing different Melitaeini host species and their relatives based on the ten concatenated mitochondrial and nuclear genes (16s, *18s, 28s, COI, EF1A1, INX, ITS2, OPS, SLD5,* and *TCEB*). The specimens are labelled by the names of the *Cotesia* species and Melitaeini host caterpillar species. Bootstrap support values (1000 replicates) are indicated for supported branches (≥60). Colors represent the main clades (A, B, C, D). Branches corresponding to the same species were collapsed for visual clarity and represented by triangles.

#### (II) Nuclear-gene dataset

The nuclear-gene dataset included 112 wasp specimens. The concatenated sequence was 2413 bp long, of which 1035 sites (43%) represented unique site-patterns, and of these 494 (20% of all sites) were parsimony informative. The model partitioning algorithm revealed five partitions in the nuclear-gene dataset (Table S2).

The ML analysis also revealed four major clades with high support (bootstrap = 100, 82, 87) and suggests that the species attacking Melitaeini butterflies are polyphyletic (Figure 2). The main clades contain the same species as the ten-gene tree, although the topology within clades A and D differ:

- **Clade A** contains the *C. acuminata agg., C. bignelli,* and *C. euphydryidis* parasitizing Melitaeini butterflies. The relationships within this clade are largely the same as in the ten-gene tree, except that *C. acuminata Sp. B* emerging from *M. scotosia* is grouped with *C. acuminata Sp. K.* Again, the relationships between the cryptic species have lower support (bootstrap = 60) (Figure 2).
- **Clade B** contains *C. congregata, C. glomerata*, and *C. koebelei* with high support (bootstrap = 95 and above) (Figure 2). Which is congruent with the phylogeny based on the ten-gene dataset.
- **Clade C** contains *C. chilonis, C. ruficrus, C. typhae,* and *C. vestalis* with high support (bootstrap = 90 and above) (Figure 2), Which is congruent to the ten-gene dataset.
- **Clade D** contains *C. melitaearum agg* and *C. cynthiae* parasitizing Melitaeini butterflies.

**Figure 2.**
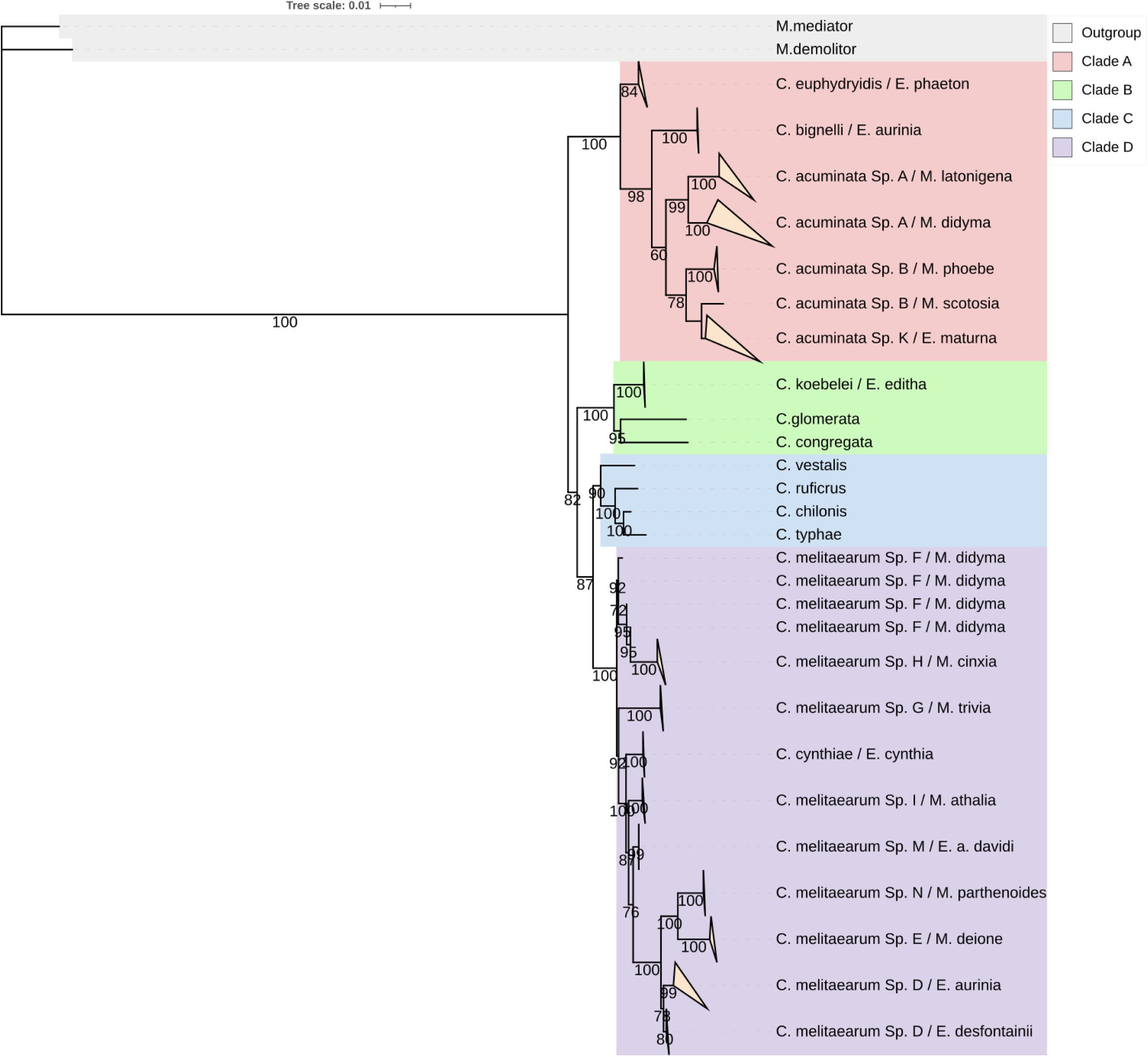
Maximum likelihood phylogeny of *Cotesia* species parasitizing different Melitaeini host species and their relatives based on the nuclear genes (*18s, 28s, EF1A1, INX, ITS2, OPS, SLD5,* and *TCEB*). The specimens are labelled by the names of the *Cotesia* species and Melitaeini host caterpillar species. Branches corresponding to the same species were collapsed for visual clarity and represented by triangles. Bootstrap support values (1000 replicates) are indicated for supported branches (≥60). Colors represent the main clades (A, B, C, D).

The relationships are moderately to strongly supported (bootstrap = 76 to 100) and show some discordance from the ten-gene phylogeny. For example, *C. melitaearum Sp. F* is grouped together with *C. melitaearum Sp. H* and is no longer polyphyletic, thus resulting in nine subclades. Additionally, *C. cynthiae* and *C. melitaearum Sp. M* are no longer sister groups. Meanwhile, the relationships between *C. melitaearum Sp. N, E* and *D* are concordant with the ten-gene dataset.

#### Mitochondrial-gene dataset

The mitochondrial-gene dataset included the sequences of the*16s* and *COI* genes from 92 specimens. The mitochondrial alignment was 1072bp long, of which 451 (42%) represented unique site-patterns, of which 242 sites (23% of all sites) were parsimony informative. The model partitioning algorithm revealed two partitions in the mitochondrial gene dataset, which correspond to the two markers (Table S3). No specimens from *C. euphydryidis* and *C. melitaearum Sp. F* from Hungary amplified for either mitochondrial marker, and no sequences were found from *C. typhae* and *C. vestalis* genomic projects, thus these species are missing from the mitochondrial phylogenetic analysis.

The ML analysis revealed the same four major clades as the ten-gene and nuclear gene phylogenies, however with varying support (bootstrap = 100, 49, 97) and topology.

- **Clade B,** which includes *C. congregata, C. glomerata* and *C. koebelei* now forms the basal clade of the phylogeny but the relationships within the clade exposed in the ten-gene and nuclear phylogenies are conserved here too.
- **Clade A** with **Clade C** and **Clade D** now group together as sister taxa to Clade B. Clades C and D are sister clades to Clade A, which is congruent the ten-gene phylogeny. The relationships within Clade A are congruent to the ten-gene phylogeny, despite missing *C. euphydryidis*. The relationships between *C. acuminata agg.* species is low (bootstrap = 60). Clade C contains *C. ruficrus* and *C.* chilonis and is congruent with the ten-gene and nuclear phylogenies, despite missing two species. Within clade D, we can still observe the paraphyletic nature of this cryptic complex, forming nine subclades comprised of the *C. melitaearum agg.* and *C. cynthiae*. Additionally, *C. melitaearum Sp. D* does not cluster by the two hosts. Many of the relationships in this clade are however unsupported (bootstrap < 60) (Figure 3).

**Figure 3.**
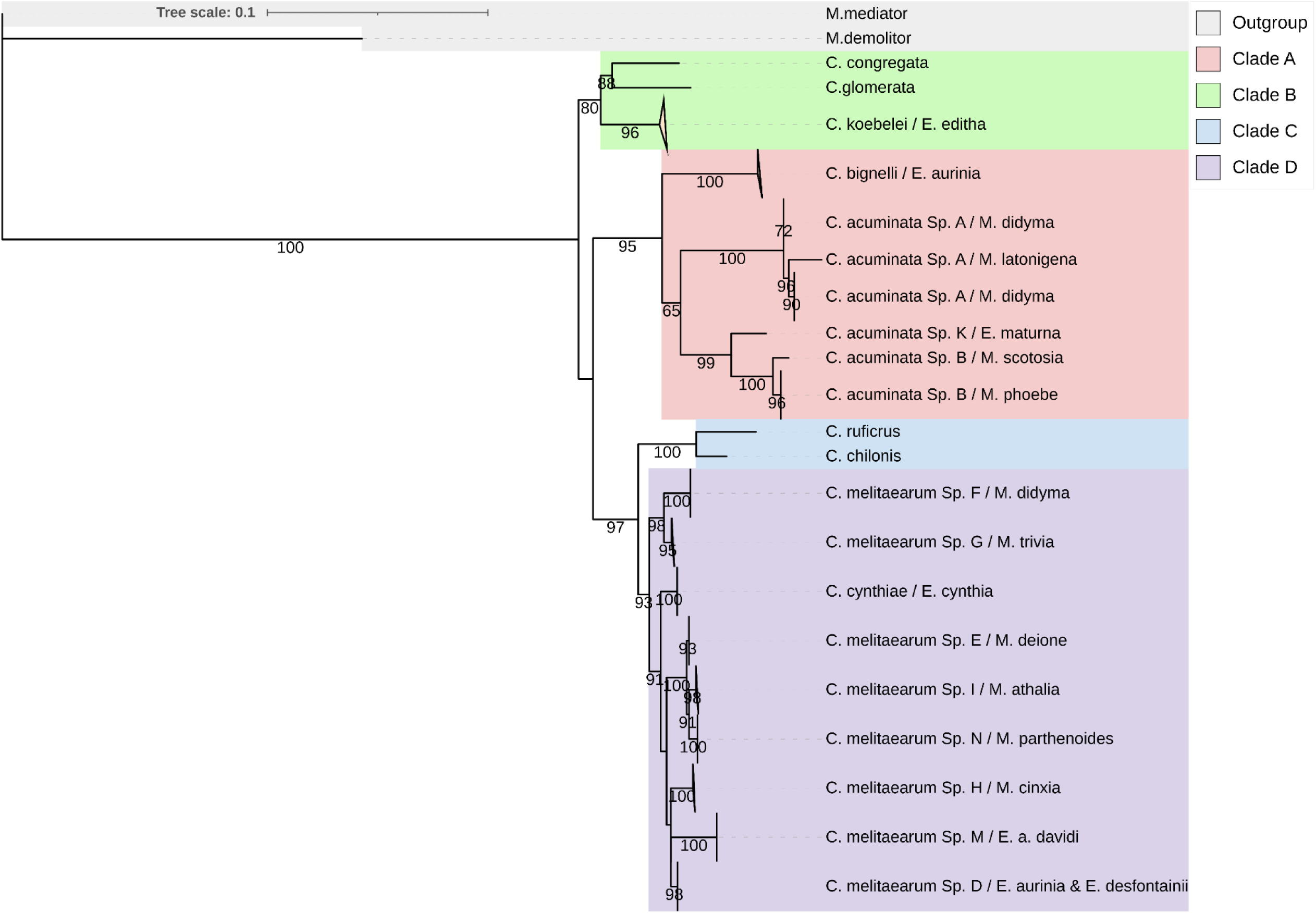
Maximum likelihood phylogeny of *Cotesia* species parasitizing different Melitaeini host species and their relatives based on the mitochondrial genes (*16s* and *COI*). The specimens are labelled by the names of the *Cotesia* species and Melitaeini host caterpillar species. Branches corresponding to the same species were collapsed for visual clarity and represented by triangles. Bootstrap support values (1000 replicates) are indicated for supported branches (≥60). Colors represent the main clades (A, B, C, D).

#### Individual Genes

All genes were also analysed individually using ML to identify discordance or outlier genes. The substitution models are listed in Table S4. Rather expectedly, the individual gene phylogenies had much lower support than either of the other phylogenies, likely because of the small nucleotide sample size included in each analysis. Alone, the individual gene trees are not phylogenetically informative. The major clades were only recovered in the *16s*, *COI*, and *SLD5* trees, but the relationships within the clades, particularly within the Clade C, remain incongruent (Figure S1-S10)

## Discussion

Resolving species phylogenies is essential for studying trait evolution, diversification, and ecological interactions, but can remain challenging in non-model cryptic species, due to limited access to sampling and molecular resources. By designing new primers and increasing the number of available molecular markers from two to ten, we were able to produce the most robust phylogeny of the *Cotesia* species parasitizing Melitaeini butterflies to date. This improved phylogeny informs on previously hidden species diversity within this genus and is an important resource to future research on Braconidae population genetics, evolutionary dynamics and ecology.

Our analyses revealed all specimen’s group in four highly supported clades (A-D), each subdivided into distinct subclades of (cryptic) species. By including genetic sequences from *Cotesia* species that do not attack Melitaeini butterflies, we confirm that the Melitaeini-specialized species are polyphyletic, appearing in three of the four main *Cotesia* clades. This suggests that *Cotesia* wasps have shifted hosts multiple times throughout their evolutionary history. Such patterns of host shifts have been observed in other parasitoid lineages, including Ichneumonoidea (Quicke 2012) and parasitoids of leaf miners (Leppänen et al. 2012) indicating that parasitoid evolution often involves ecological switching rather than strict co divergence with hosts.

Many of these Melitaeini-associated *Cotesia* species occur in sympatry, yet the drivers of radiation and barriers to gene flow remain unknown (Kankare et al. 2005b, Valerio et al. 2023). Host evolution appears to play a partial role: for example, in Clade A, the basal species parasitize *Euphydryas* caterpillars, while most *C. acuminata* species have shifted to *Melitaea.* Given that *Euphydryas* butterflies diverged before *Melitaea* (Long et al. 2014), it likely supports the hypothesis that the ancestral host for Clade A belonged to the *Euphydryas* butterfly lineage. A similar pattern may exist in Clade B where *C. koebelei* parasitizes *E. editha* and has also been tentatively reported from *Melitaea* (Moore 1989). If *C. koebelei* parasitizes both *Euphydryas* and *Melitaeini* butterflies, it could represent a cryptic species complex or a rare generalist among these species able to attack both genera, emphasizing the need for further sampling of North American Melitaeini butterflies and their *Cotesia* parasitoids.

In contrast, Clade D shows an opposite pattern to Clade A: with basal species parasitizing *Melitaea*, followed by a shift to *Euphydryas*. This may reflect a retained ability to use *Euphydryas* inherited from a shared ancestor with Clade A, possibly representing a recolonization event. Species from Clade C are not known to use Melitaeini butterflies but span a wider range of Lepidoptera host taxa. For example, *C. vestalis* parasitizes *Plutella xylostella* (Yponomeutoidea), a lineage near the base of the Lepidopteran tree (Kawahara et al. 2019), while more recently diverged *Cotesia* in this clade parasitize more recently diverged groups such as Pyraloidea and Noctoidae.

Although the relationships within and between cryptic species were better supported in our final phylogeny compared to previous studies (Kankare and Shaw 2004), comparisons of the tree topologies suggested conflicts between nuclear and mitochondrial reconstructions. The differences between analyses might be due to the non-random absence of mitochondrial data for several taxa in two clades, which can bias phylogenetic inference, especially when relatively few markers are used (Xi et al. 2016). In addition, the discordance within clades may reflect geneflow between lineages such as hybridization or incomplete lineage sorting (Zadra et al. 2021).

While we bring an improved version of the *Cotesia* phylogeny, our data remains too weak to fully evaluate the level of hybridization and introgression between the lineages, or the frequency of host shifts within clades and could have mislead phylogenetic inference especially under incomplete lineage sorting (Xi et al. 2016). In the future, the use of additional genetic material, specimens from other species, or a wider geographic range will be needed to confidently evaluate the extent of the true evolutionary history of these events.

Such increased sampling would allow for explicit research on the contributions of geography, ecology, and host associations in the diversification of *Cotesia*. For example, other parasitoid lineages show patterns of latitudinal diversity gradients and geographic structuring (Castellanos-Labarcena et al. 2025) and similarities might be revealed in *Cotesia*. Additionally, the butterfly host ecology could be explored, which has been observed in parasitoids of leaf miners that often radiate to unrelated hosts that share the same host plant (Leppänen et al. 2012). Finally studying host associations and range may also influence *Cotesia* diversity through mechanisms such as host fidelity. Whereby many parasitoid wasp females preferentially oviposit on the same host species from which they emerged, reinforcing reproductive barriers and potentially promoting speciation (Kimura and Novković 2015, Hood et al. 2015, Forbes et al. 2017). Network based approaches provide promising tools to study host-parasitoid coevolution (Braga et al. 2018), testing hypotheses such as adaptive radiation (Ehrlich and Raven 1964), or host use oscillations (Janz and Nylin 2008). Understanding host repertoire has important implications for conservation, as species with narrow fundamental host ranges face greater extinction risks, and identifying such species can help prioritize conservation efforts (Farrell et al. 2021).

## Acknowledgements

Thanks to the members of the ISEE lab and life-history evolution group at the University of Helsinki for the many discussions and input. Thank you to Victoria Twort for her input on the manuscript.

Thanks to everyone involved in the sample collection, including Constantí Stefanescu and his team in Spain, Yannick Chittaro in Switzerland, and in Finland Suvi Ikonen and Krista Raveala, along with those involved in the Åland butterfly survey.

## Data Availability

Data and material is available from the authors upon request.

## Notes

### Competing Interest Statement

The authors have declared no competing interest.

### Funding

- Research Council of Finland: 328944

