## Supplementary Materials for "Progress in the molecular Phylogeny of *Cotesia acuminata* and *C. melitaearum* cryptic species complexes"

Table S1. Ten-gene dataset partitions determined by partitioning algorithm, and best fit model as determined by ModelFinder using BIC.

| **Partition ID** | **Genes** | **Model** | **BIC** |
| --- | --- | --- | --- |
| 1 | *16s, ITS2* | K3Pu+F+G4 | 8136.897 |
| 2 | *18s, 28s* | HKY+F+R2 | 5778.127 |
| 3 | *COI, INX, SLD5* | TIM+F+I+R2 | 10450.674 |
| 4 | *EF1a1* | GTR+F+I+G4 | 2328.192 |
| 5 | *OPS, TCEB* | K3Pu+F+I+R2 | 3207.386 |

Table S2. Nuclear gene dataset partitions determined by partitioning algorithm, and best fit model determined by ModelFinder using BIC.

| **Partition ID** | **Genes** | **Model** | **BIC** |
| --- | --- | --- | --- |
| 1 | *18s, 28s* | HKY+F+R2 | 5909.165 |
| 2 | *EF1a1* | GTR+F+I+G4 | 2351.312 |
| 3 | *INX, SLD5* | HKY+F+G4 | 4031.702 |
| 4 | *ITS2* | GTR+F+I+G4 | 4139.166 |
| 5 | *OPS, TCEB* | K3Pu+F+I+R2 | 3202.952 |

Table S3. Mitochondrial-gene dataset partitions determined by partitioning algorithm, and best fit model determined by ModelFinder using BIC.

| **Partition ID** | **Genes** | **Model** | **BIC** |
| --- | --- | --- | --- |
| 1 | *16s* | K3Pu+F+G4 | 3421.950 |
| 2 | *COI* | TIM+F+G4 | 5842.849 |

Table S4. Substitution models used for individual genes, determined by ModelFinder, and selected based on BIC.

| **Gene** | **Model** | **BIC** |
| --- | --- | --- |
| 16s | K3Pu+F+G4 | 4508.990 |
| 18s | K2P+G4 | 2307.056 |
| 28s | HKY+F+R2 | 4817.051 |
| COI | TIM+F+G4 | 6990.912 |
| EF1A1 | TIM2e+I | 3155.489 |
| INX | HKY+F+G4 | 2515.397 |
| ITS2 | F81+F+G4 | 4881.005 |
| OPS | TPM2u+F+G4 | 1940.479 |
| SLD5 | K3P+G4 | 2590.048 |
| TCEB | HKY+F+G4 | 2169.188 |

Table S5. Characteristics of the additional primer pairs created for this study. Includes the marker targeted (gene), primer name, primer sequence, amplicon size. “-” indicates the information was unavailable.

| **Gene** | **Primer name** | **Sequence (5'-3')** | **Amplicon size** |
| --- | --- | --- | --- |
| *DAP1* | DAP-F | 5`-CAAAACACCACGGGACGAAC | 132 |
|  | DAP-R | 5`-TCTGGAGGAAAATCGGCGTT |  |
| *PSMB* | PSMB-F | CGCCTGAGATTCCGGCTAAA | 368 |
|  | PSMB-R | 5`-TGTACGATAGCACCCCAACC |  |
| *SBDS* | SBDS-F | 5`-TCGGCAATGACAACCAGACA | 305 |
|  | SBDS-R | 5`-GACCCTAACTCGCATCCTGG |  |
| *NUTF2* | NUTF2-F | 5`-GCTCAATAGCCTGGGTTTCCA | 256 |
|  | NUTF2-R | 5`-CAGTGGCTTCAGCACAAACG |  |
| *CPSF5* | CPSF5-F | 5`-GCGTAGGAGTGTCGAAGGAG | 375 |
|  | CPSF5-R | 5`-AGTTCAAACAATGGCGCAGC |  |
| *HIGD2A* | HIGD2A-F | 5`-TCCTTTGGTGCCGATAGGTG | 128 |
|  | HIGD2A-R | 5`-TGAAACCCTGAGCAGATACCC |  |
| *AIMP1* | AIMP1-F | 5`-AGCTCTGGGATGACGTTGAC | 329 |
|  | AIMP1-R | 5`-TTTTGGCGAGTTTTGGTGCG |  |
| *ATG3* | ATG3-F | 5`-ATGGGCAACTGGAGACGAAG | 724 |
|  | ATG3-R | 5`-CGGAATCACCGACTGGACAA |  |
| *INTS2* | INTS2-F | 5`-GGTACAGAAGTCGGTGAGGC | 309 |
|  | INTS2-R | 5`-CAGCAGCGGCGAGAATAAAC |  |
| *Taz* | Taz-F | 5`-TTCGCAAGAAAACGACGCAC | 371 |
|  | Taz-R | 5`-TCATGTGCTGCTAACGACCA |  |
| *B4GALT7* | B4GALT7-F | 5`-CAGTTTGCTCCGCACATGAAA | 357 |
|  | B4GALT7-R | 5`-ATCTTCTAAGCCCCAACCCC |  |
| *SMD2* | SMD2-F | 5`-GGACCACTTTCAGTACTTACCCA | 273 |
|  | SMD2-R | 5`-GGCGGTTGCAAGTGGATTTC |  |
| *STX6* | STX6-F | 5`-CGTGACAGAACTGCGAGACA | 388 |
|  | STX6-R | 5`-ATGCAGTTGTAGGAGCAGGA |  |

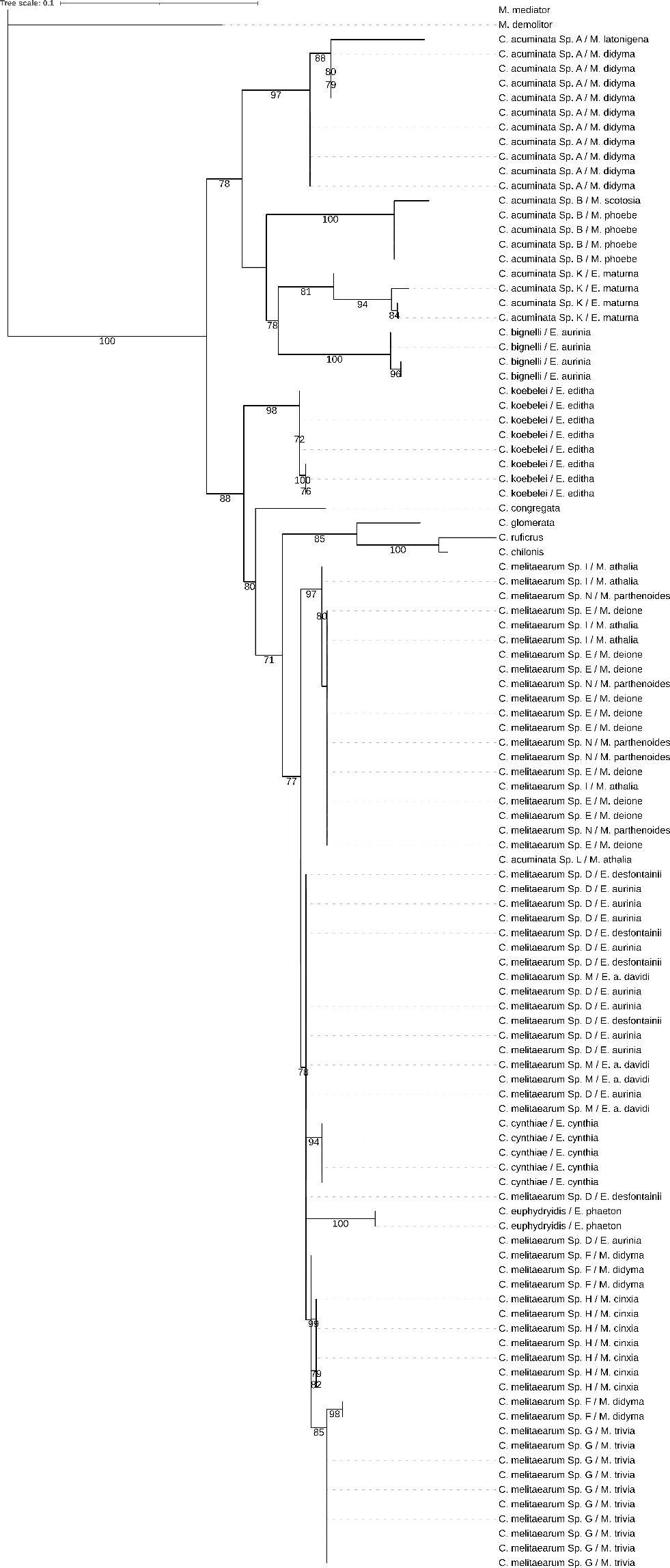

Figure S1. Maximum likelihood phylogeny of *Cotesia* species parasitizing different Melitaeini host species and their relatives based on the mitochondrial gene *16s.* The specimens are labelled by the names of the *Cotesia* species and Melitaeini host caterpillar species*.* Bootstrap support values (1000 replicates) are indicated for supported branches (≥60).

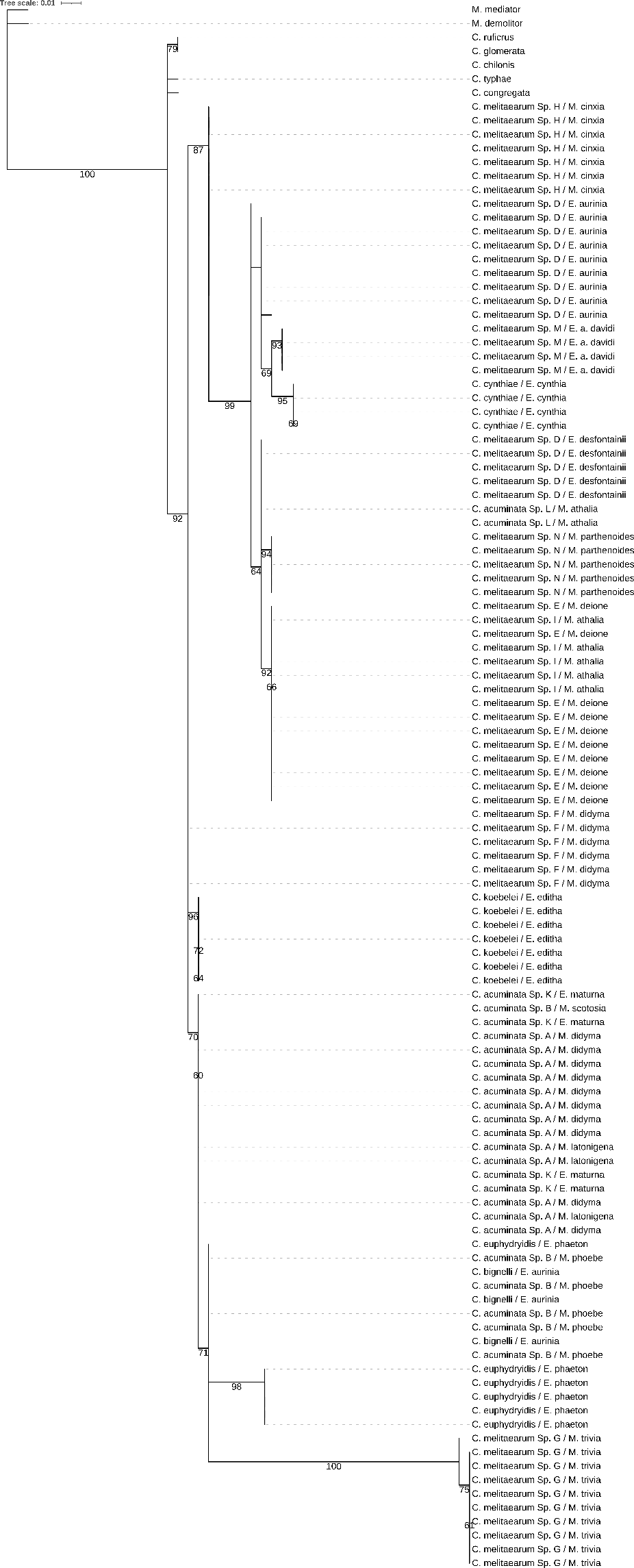

Figure S2. Maximum likelihood phylogeny of *Cotesia* species parasitizing different Melitaeini host species and their relatives based on the nuclear gene *18s.* The specimens are labelled by the names of the *Cotesia* species and Melitaeini host caterpillar species. Bootstrap support values (1000 replicates) are indicated for supported branches (≥60).

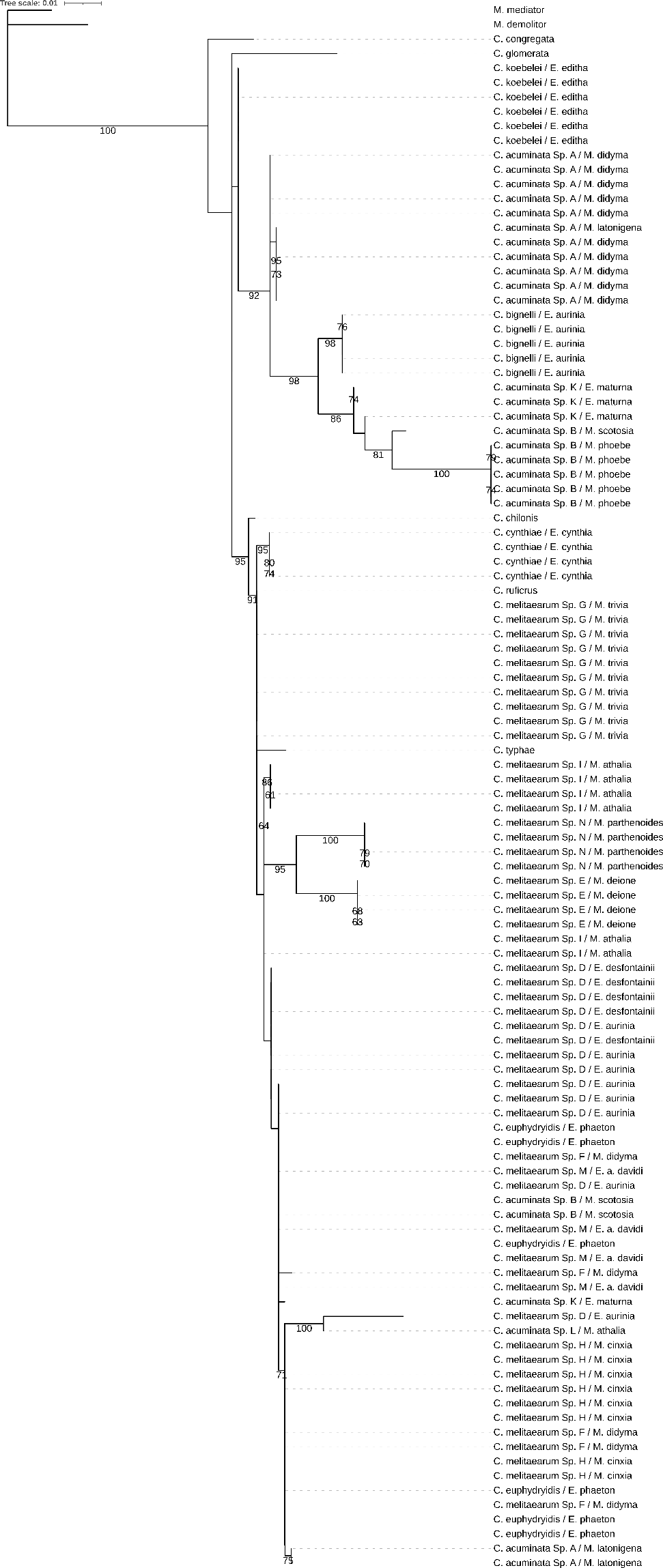

Figure S3. Maximum likelihood phylogeny of *Cotesia* species parasitizing different Melitaeini host species and their relatives based on the nuclear gene *28s.* The specimens are labelled by the names of the *Cotesia* species and Melitaeini host caterpillar species. Bootstrap support values (1000 replicates) are indicated for supported branches (≥60).

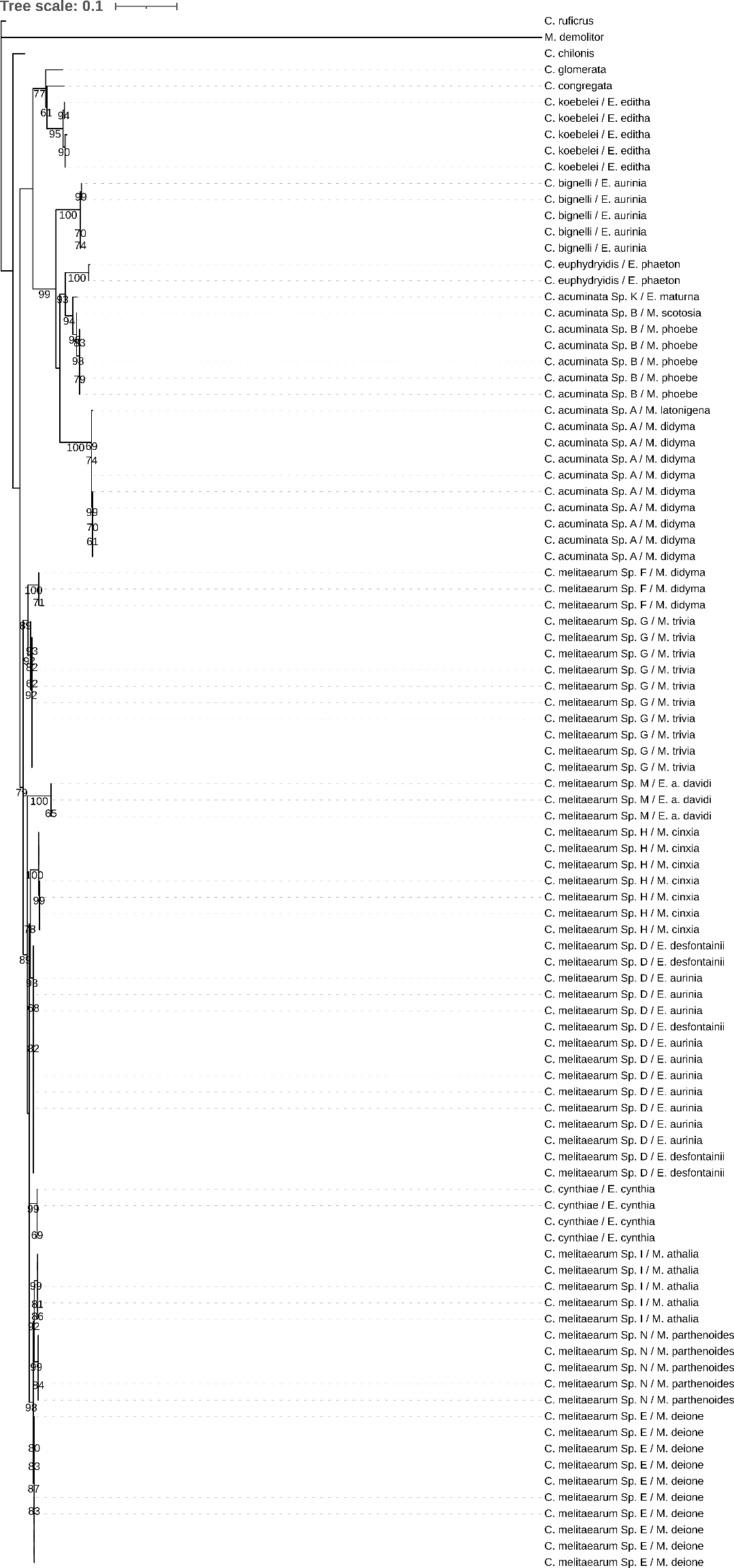

Figure S4. Maximum likelihood phylogeny of *Cotesia* species parasitizing different Melitaeini host species and their relatives based on the mitochondrial gene *COI.* The specimens are labelled by the names of the *Cotesia* species and Melitaeini host caterpillar species. Bootstrap support values (1000 replicates) are indicated for supported branches (≥60).

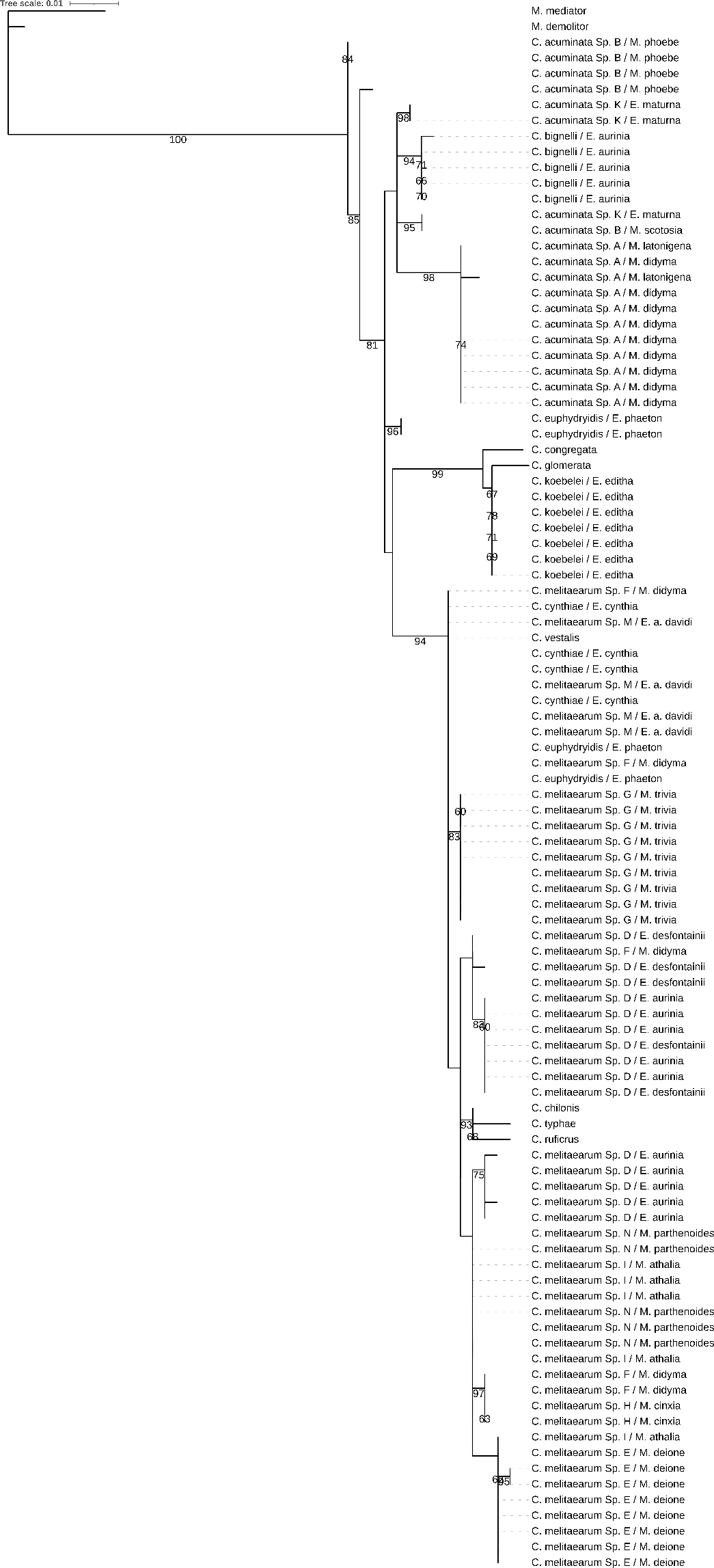

Figure S5. Maximum likelihood phylogeny of *Cotesia* species parasitizing different Melitaeini host species and their relatives based on the nuclear gene *EF1a1.* The specimens are labelled by the names of the *Cotesia* species and Melitaeini host caterpillar species. Bootstrap support values (1000 replicates) are indicated for supported branches (≥60).

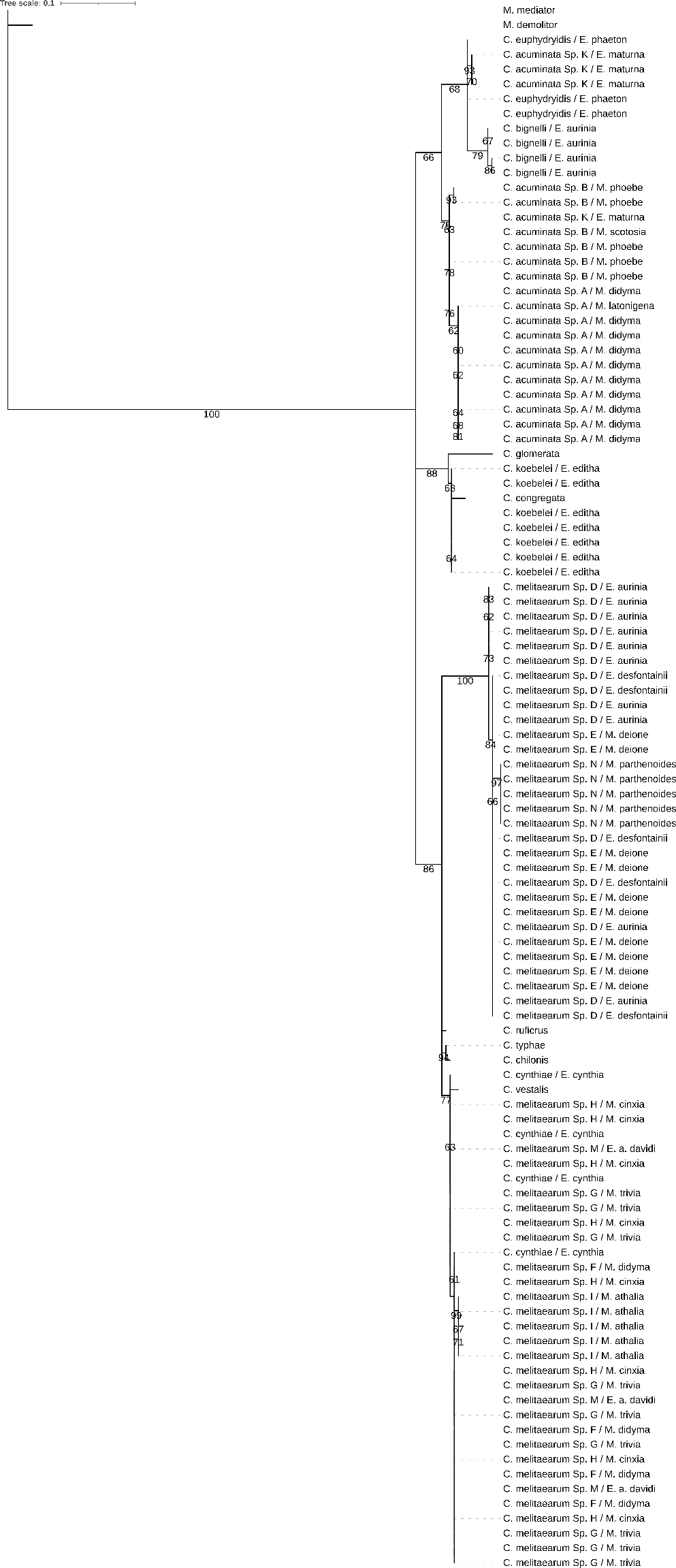

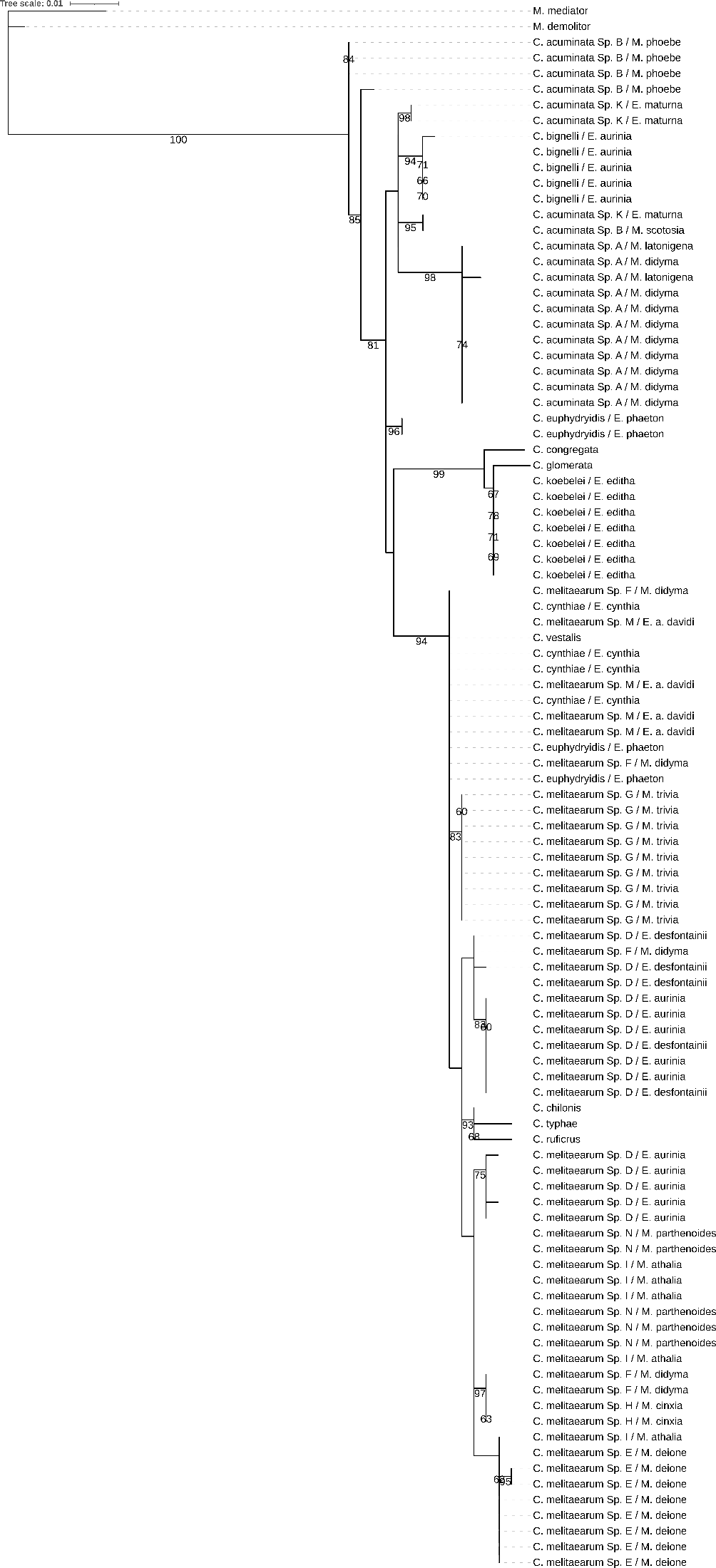

Figure S6. Maximum likelihood phylogeny of *Cotesia* species parasitizing different Melitaeini host species and their relatives based on the nuclear gene *INX.* The specimens are labelled by the names of the *Cotesia* species and Melitaeini host caterpillar species. Bootstrap support values (1000 replicates) are indicated for supported branches (≥60).

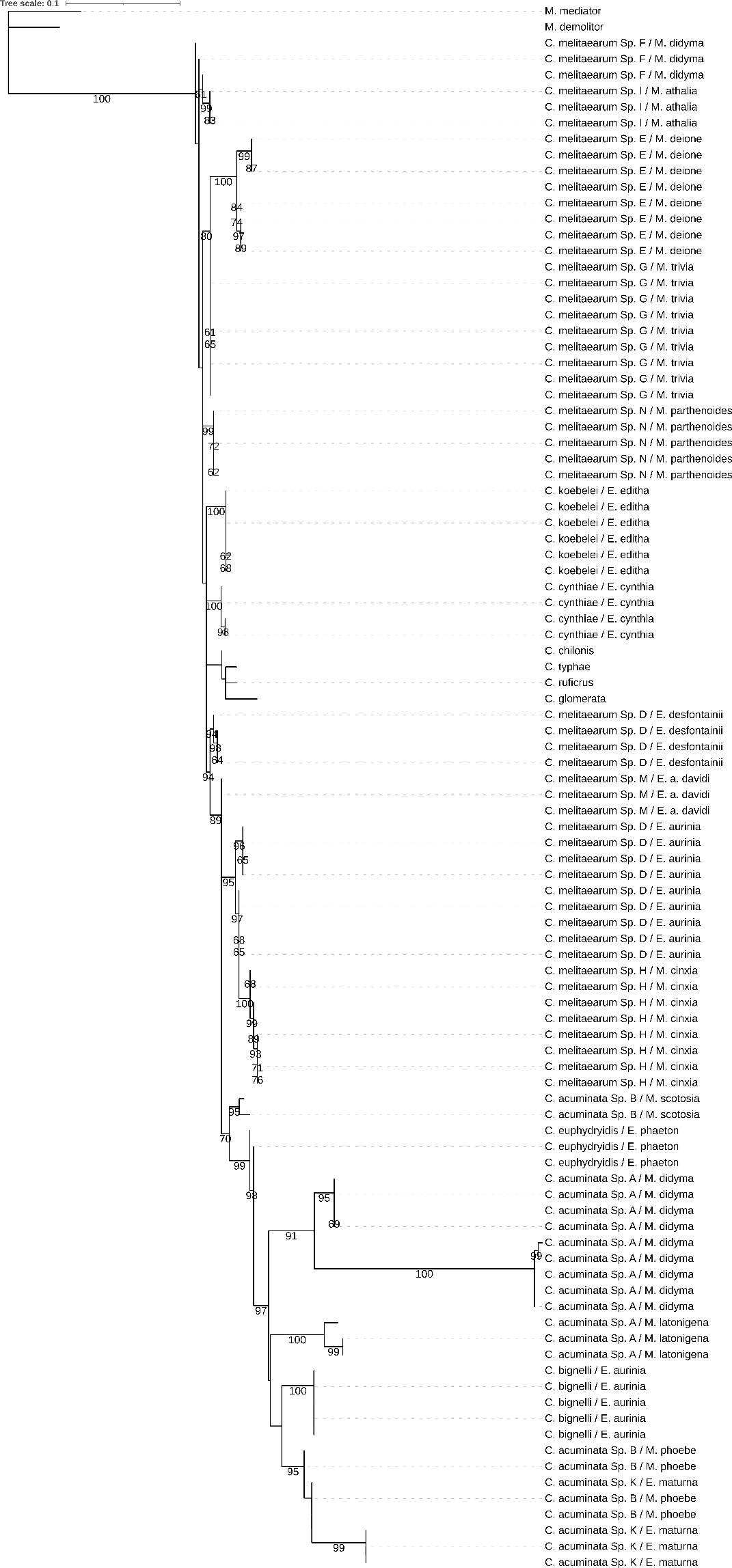

Figure S7. Maximum likelihood phylogeny of *Cotesia* species parasitizing different Melitaeini host species and their relatives based on the nuclear gene *ITS2.* The specimens are labelled by the names of the *Cotesia* species and Melitaeini host caterpillar species. Bootstrap support values (1000 replicates) are indicated for supported branches (≥60).

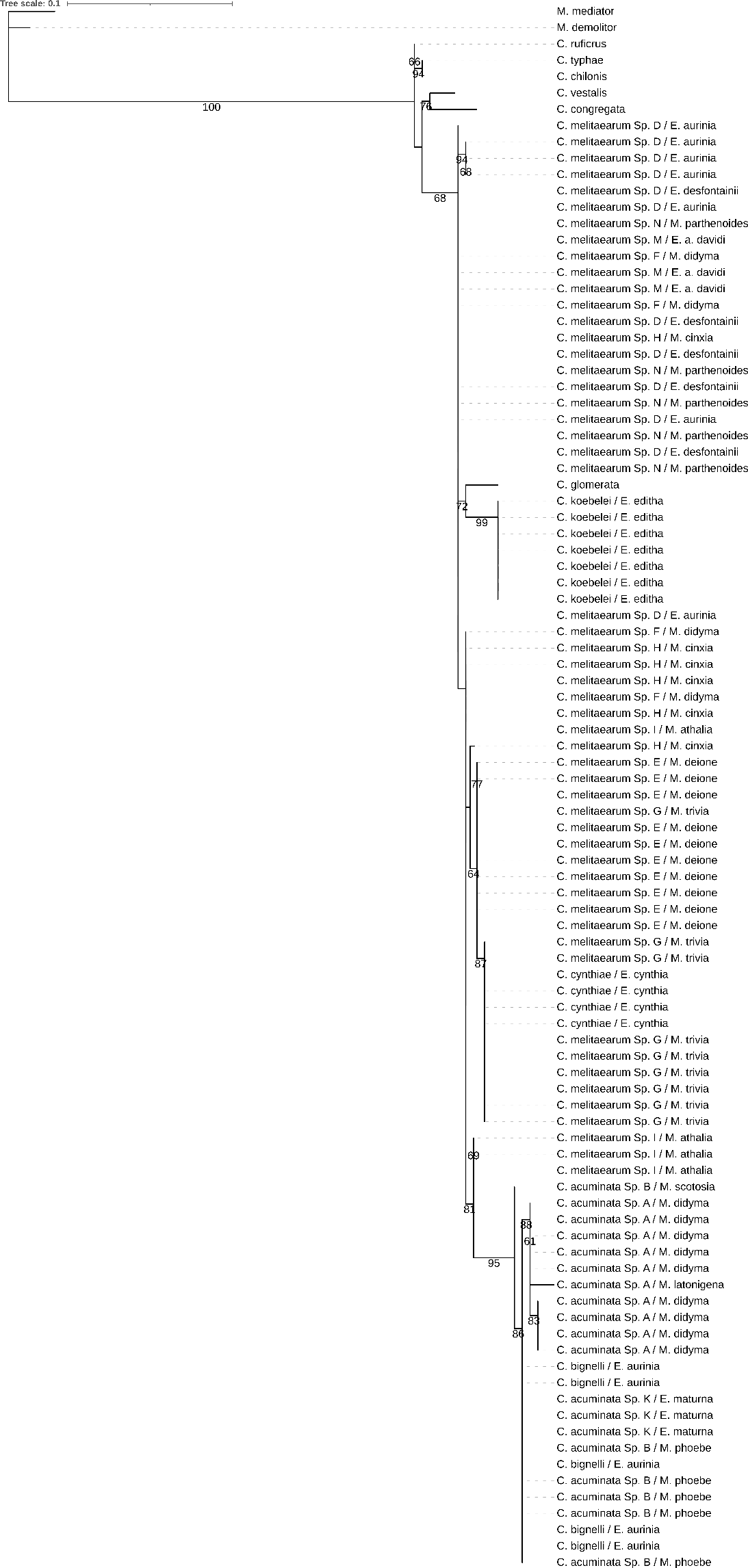

Figure S8. Maximum likelihood phylogeny of *Cotesia* species parasitizing different Melitaeini host species and their relatives based on the nuclear gene *Ops.* The specimens are labelled by the names of the *Cotesia* species and Melitaeini host caterpillar species. Bootstrap support values (1000 replicates) are indicated for supported branches (≥60).

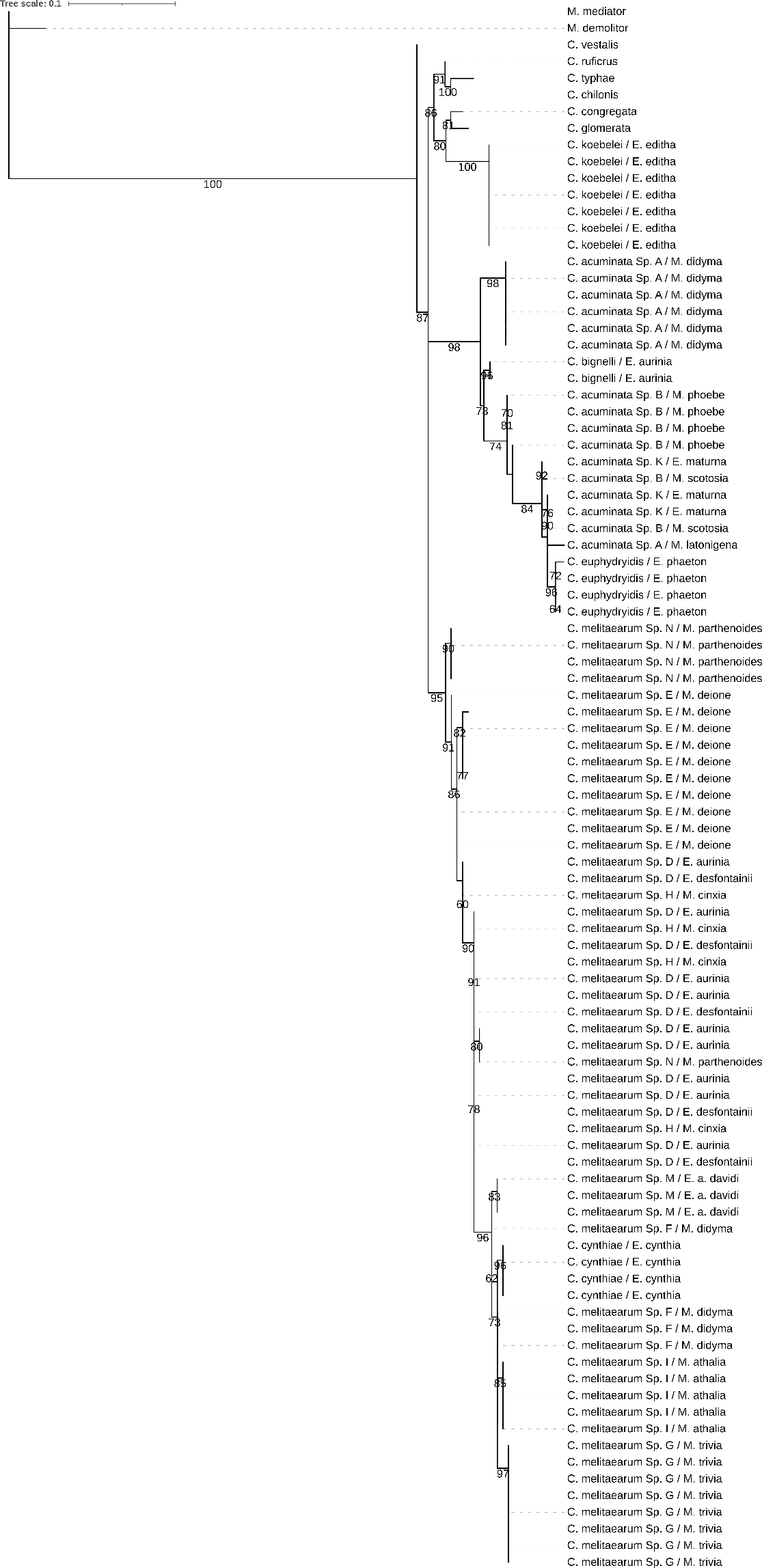

Figure S9. Maximum likelihood phylogeny of *Cotesia* species parasitizing different Melitaeini host species and their relatives based on the nuclear gene *SLD5.* The specimens are labelled by the names of the *Cotesia* species and Melitaeini host caterpillar species. Bootstrap support values (1000 replicates) are indicated for supported branches (≥60).

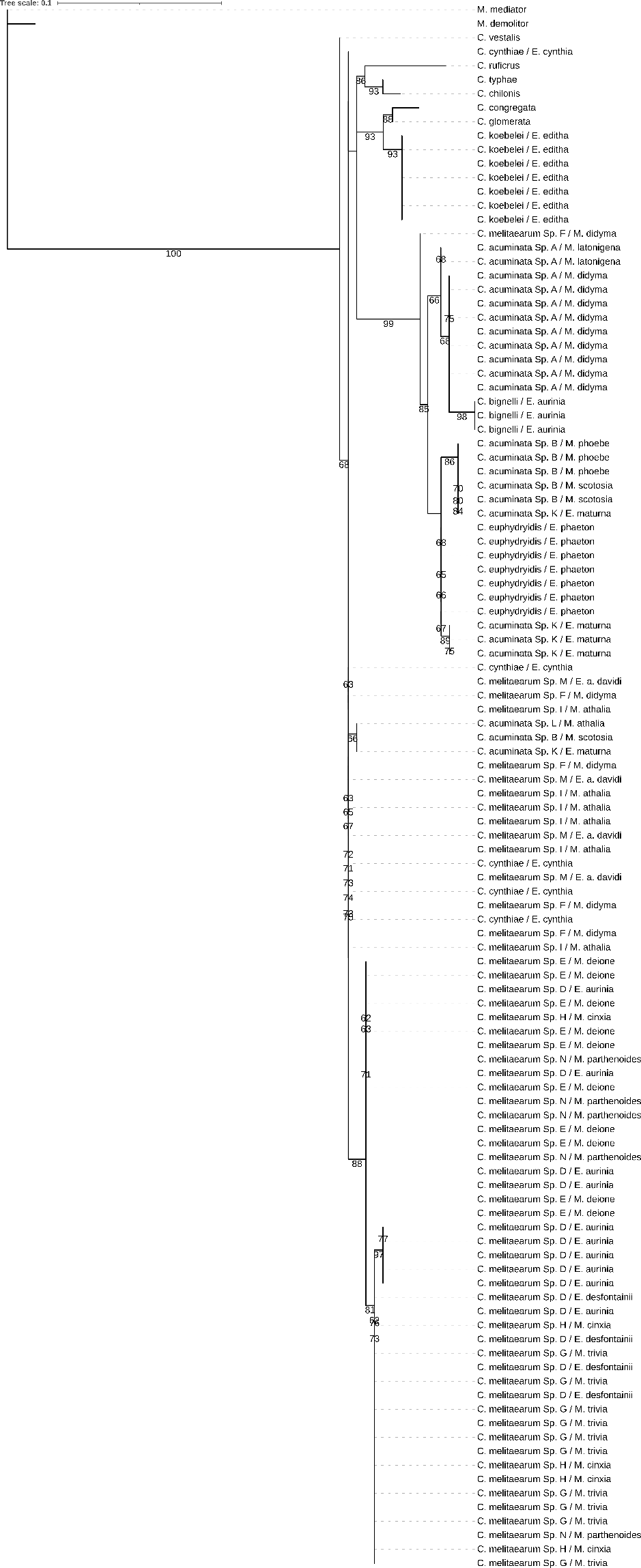
TCEB

Figure S10. Maximum likelihood phylogeny of *Cotesia* species parasitizing different Melitaeini host species and their relatives based on the nuclear gene *TCEB.* The specimens are labelled by the names of the *Cotesia* species and Melitaeini host caterpillar species. Bootstrap support values (1000 replicates) are indicated for supported branches (≥60).
